# Identifying indels from WGS short reads of haploid genomes distinguishes variant-calling algorithms

**DOI:** 10.1101/2022.12.14.520524

**Authors:** Yee Mey Seah, Mary K. Stewart, Daniel Hoogestraat, Molly Ryder, Brad T. Cookson, Stephen J. Salipante, Noah G. Hoffman

## Abstract

Identification of clinically relevant strains of bacteria increasingly relies on whole genome sequencing. The downstream bioinformatics steps necessary for calling variants from short read sequences are well-established but seldom validated against haploid genomes. We devised an *in silico* workflow to introduce single nucleotide polymorphisms (SNP) and indels into bacterial reference genomes, and computationally generate sequencing reads based on the mutated genomes. We then applied the method to *Mycobacterium tuberculosis* H37Rv and used the synthetic reads as truth sets for evaluating several popular variant callers. Insertions proved especially challenging for most variant callers to correctly identify, relative to deletions and single nucleotide polymorphisms. With adequate read depth, however, variant callers that use high quality soft-clipped reads and base mismatches to perform local realignment consistently had the highest precision and recall in identifying medium-length insertions and deletions.

## Introduction

Whole genome sequencing and identification of bacteria can be immensely useful in tracking the transmission and evolution of pathogens during outbreaks, as well as in predicting clinically relevant phenotypes such as antimicrobial resistance. Bioinformatics workflows rely on the accuracy of variant calling algorithms and pipelines to identify relevant strains and species. Accuracy is especially critical in informing clinical decisions; however, the performance of many variant callers used in clinical microbiology workflows have primarily been developed for and evaluated against human reference genomes. Truth sets of bacterial species variants are limited (Bush et al. (1)). In addition, the conclusions of variant caller validation against linear and diploid reference genomes do not always apply to variant calling in haploid and circular bacterial genomes. For example, genotyping alleles relies on setting the appropriate minimum threshold for variant allele frequencies (VAF), and the VAF threshold for genotyping homozygous versus heterozygous alleles in diploid genomes necessarily differs from the minimum threshold for identifying alternate alleles in haploid genomes; variant callers that do not account for this may not be appropriate for bacterial variant calling. Additionally, relying on aligners that do not recognize ambiguously mapped reads as potential signals of circular genomes may result in reduced coverage across linearized breakpoints.

Given these issues, various efforts to develop recommendations for benchmarking variant caller performance against bacterial reference genomes have been made, although these analyses have focused on the identification of single nucleotide variants exclusively (1–3). Bush et al. (1) evaluated bacterial SNP-calling pipelines using real and simulated reads of several species of *Enterobacteriaceae* and found that reference genome selection significantly impacts the performance of variant-calling pipelines, especially for highly recombinogenic bacterial species. This corresponds to the findings by (4), which concluded that selection of both reference genomes and short read aligners affect variant calling in the clonal *Listeria monocytogenes* species.

One reason benchmarking analyses are restricted to evaluating SNP calls is the diversity of indel-calling algorithms (1), which prohibits direct performance comparisons. Indel-containing reads are challenging to map to unique and correct genomic locations; both insertions and deletions can generate alternative haplotypes that correspond to multiple loci on reference haplotypes (5). Independent mapping of individual fragments by read mappers is also more likely to tolerate mismatches than gapped alignments (5). Therefore, in addition to alignment-based methods, other algorithms have been developed specifically for indel-calling including split read mapping, paired-end read mapping, and haplotype-based methods (6). Alignment-based indel callers rely on different models to distinguish true indels from alignment errors. Meanwhile, paired reads are used by split read mapping methods that identify discordant pairs for de novo assembly, and paired-end read mapping methods that compare expected to actual mapped distance between pairs (6). Finally, haplotype-based methods identify active regions with evidence of indels relative to a reference sequence, followed by reassembly of the active regions to generate possible haplotypes from the reads; indels are then called based on the posterior probabilities of reads realigned to the possible haplotypes.

We expand on previous efforts to validate bacterial variant-calling by developing a toolkit for introducing synthetic variants into the *Mycobacterium tuberculosis* H37Rv reference genome, and measuring the precision and recall of different callers in identifying not only SNPs, but also small insertions and deletions. We used these *in silico* altered genomes to test the performance of seven variant callers: bcftools, DeepVariant, DiscoSNP, FreeBayes, GATK HaplotypeCaller, Lancet, and VarDict (Java implementation). These variant callers are widely used, have good documentation, and appear to be actively maintained within the last five years. Algorithms for identifying sequence variants from short reads can be broadly categorized into reference-based methods that rely on mapping reads onto a curated reference genome, and newer reference-free methods that construct deBruijn graphs of k-mers; some methods are a hybrid of both. All the variant callers we tested were reference-based methods, except for DiscoSNP; Table 1 summarizes the general features of the tested variant callers.

**Table 1:**
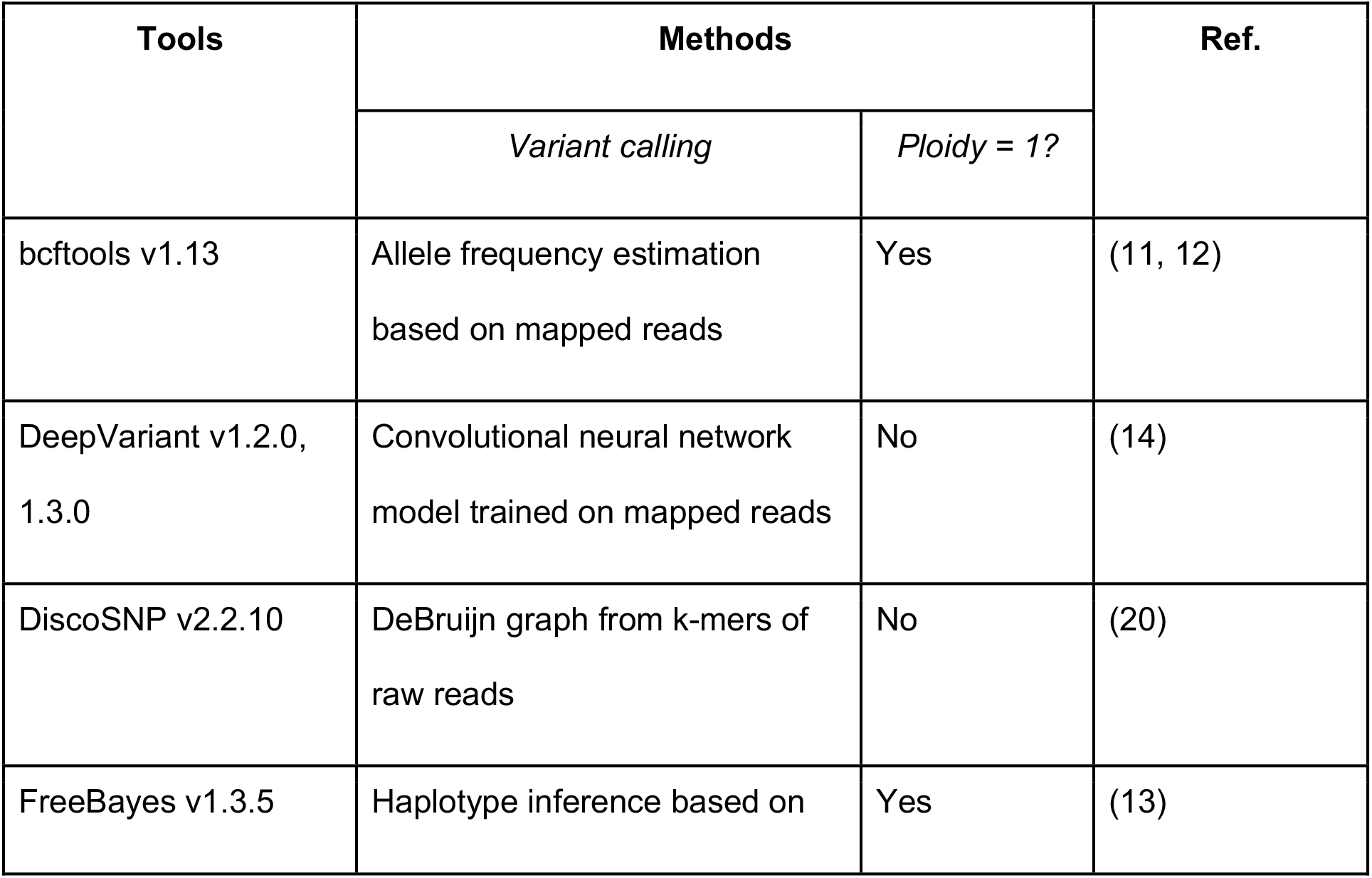
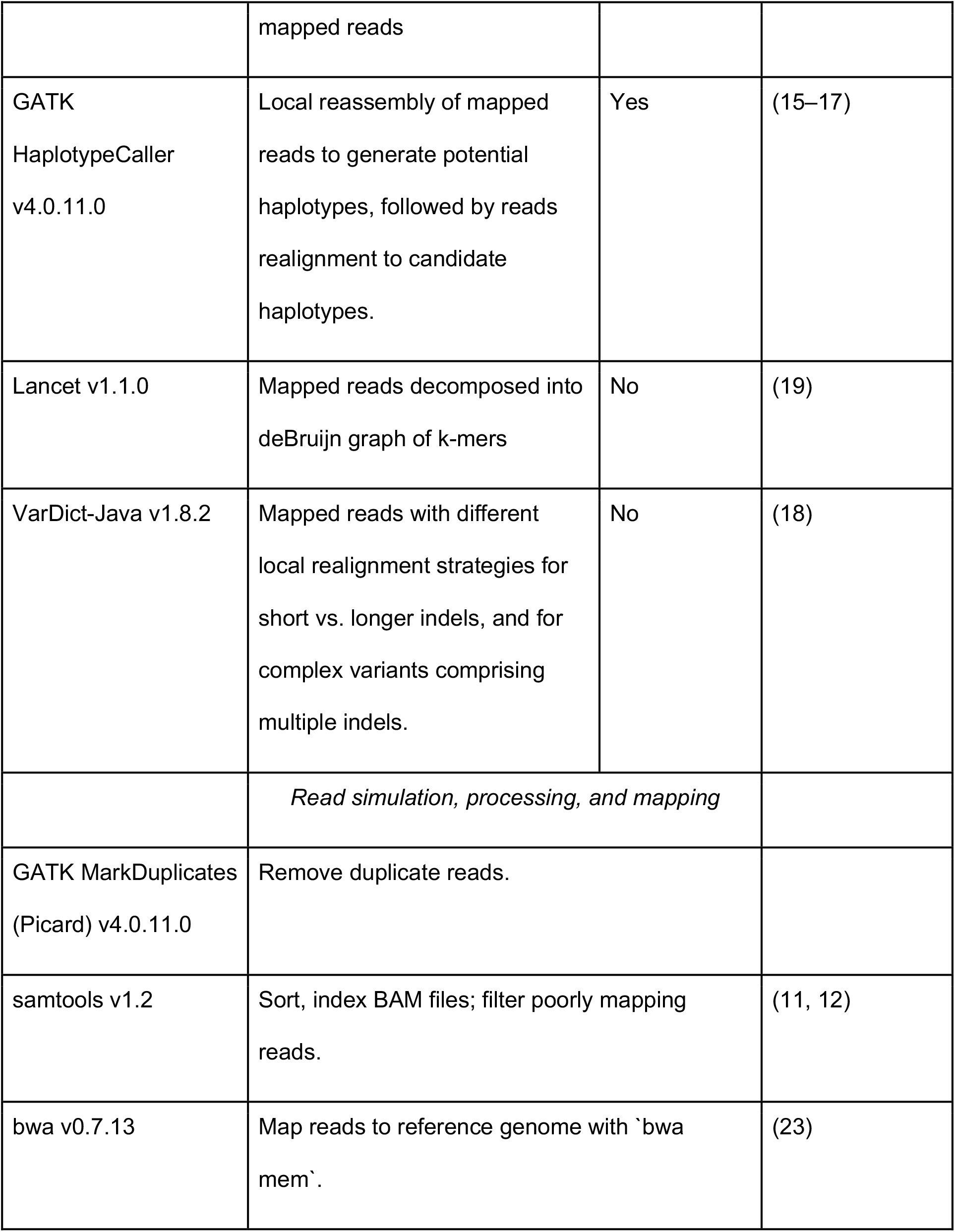
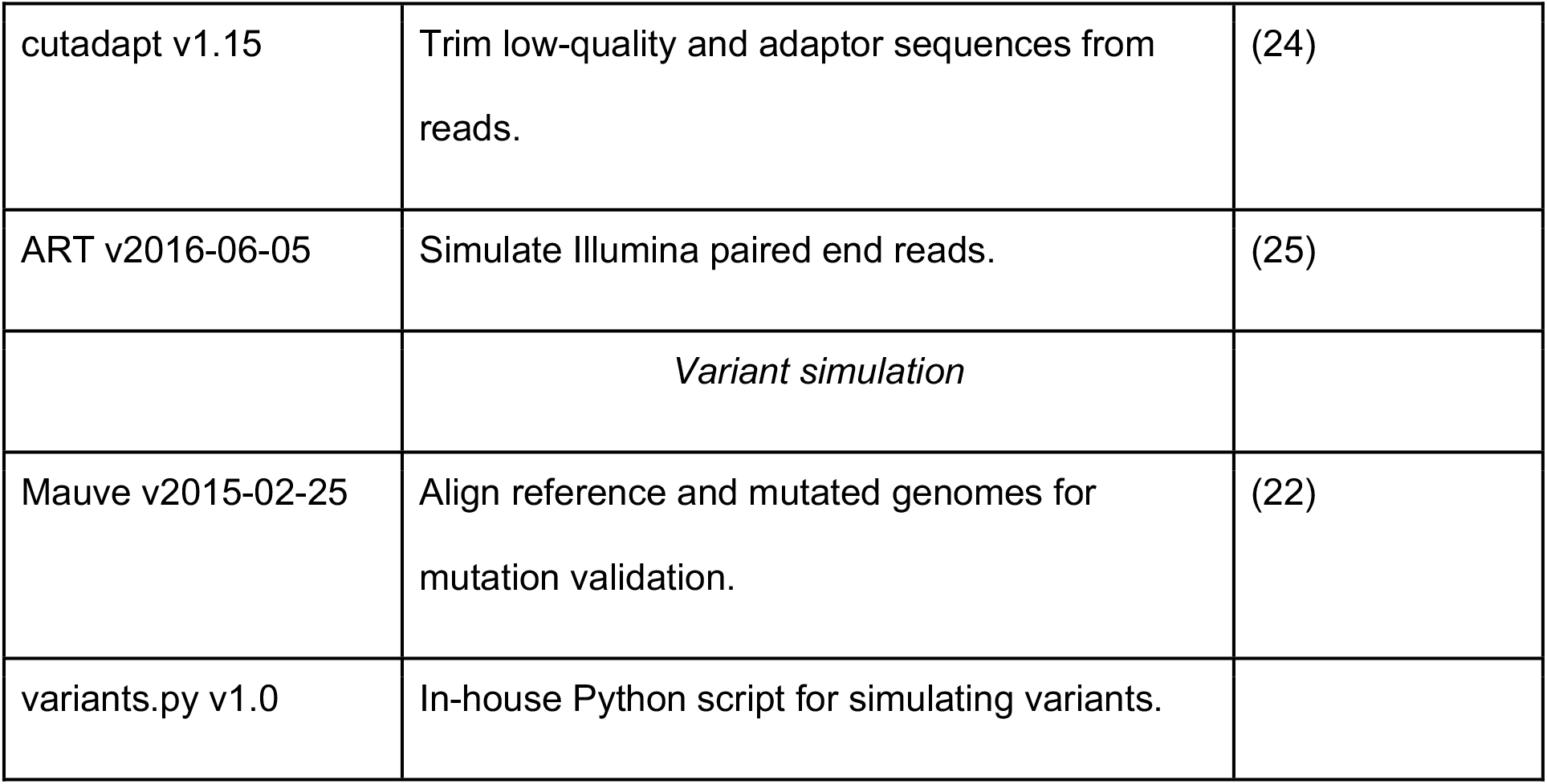
Summary of tools.

| Tools | Methods |  | Ref. |
| --- | --- | --- | --- |
|  | *Variant calling* | *Ploidy = 1?* |  |
| bcftools v1.13 | Allele frequency estimation based on mapped reads | Yes | (11, 12) |
| DeepVariant v1.2.0, 1.3.0 | Convolutional neural network model trained on mapped reads | No | (14) |
| DiscoSNP v2.2.10 | DeBruijn graph from k-mers of raw reads | No | (20) |
| FreeBayes v1.3.5 | Haplotype inference based on mapped reads | Yes | (13) |
| GATK<br>HaplotypeCaller<br>v4.0.11.0 | Local reassembly of mapped reads to generate potential haplotypes, followed by reads realignment to candidate haplotypes. | Yes | (15–17) |
| Lancet v1.1.0 | Mapped reads decomposed into deBruijn graph of k-mers | No | (19) |
| VarDict-Java v1.8.2 | Mapped reads with different local realignment strategies for short vs. longer indels, and for complex variants comprising multiple indels. | No | (18) |
|  | *Read simulation, processing, and mapping* |  |  |
| GATK MarkDuplicates<br>(Picard) v4.0.11.0 | Remove duplicate reads. |  |  |
| samtools v1.2 | Sort, index BAM files; filter poorly mapping reads. |  | (11, 12) |
| bwa v0.7.13 | Map reads to reference genome with `bwa mem`. |  | (23) |

|  |  |  |
| --- | --- | --- |
| cutadapt v1.15 | Trim low-quality and adaptor sequences from reads. | (24) |
| ART v2016-06-05 | Simulate Illumina paired end reads. | (25) |
|  | *Variant simulation* |  |
| Mauve v2015-02-25 | Align reference and mutated genomes for mutation validation. | (22) |
| variants.py v1.0 | In-house Python script for simulating variants. |  |

## Results

Sixty mutated genomes were generated *in silico* in each of the SNPs-only, insertions-only, and deletions-only datasets, resulting in approximately 1.2 million non-duplicate mutations per dataset. All seven variant callers performed similarly well in calling variants from the dataset that comprised only SNPs (Fig. 1a), with recall ranging from 97.9% (DiscoSNP) - 99.9% (bcftools, DeepVariant, FreeBayes, GATK, and VarDict), and precision 99.96% (DiscoSNP) - 100% (bcftools, FreeBayes, GATK, VarDict). Performance predictably declined when calling indels, with the most variable performance on the insertions-only dataset (Figs. 1b, 1c). Recall on the insertions-only dataset ranged from 33.29% (DiscoSNP) - 99.86% (GATK), while recall ranged from 71.20% (DiscoSNP) - 99.99% (GATK) on the datasets that consisted of only deletions. Meanwhile, precision on the insertions-only dataset ranged from 83.54% (VarDict) - 99.96% (GATK), and 97.39% (bcftools) - 99.99% (GATK and Lancet), on the deletions-only dataset.

**Figure 1.**
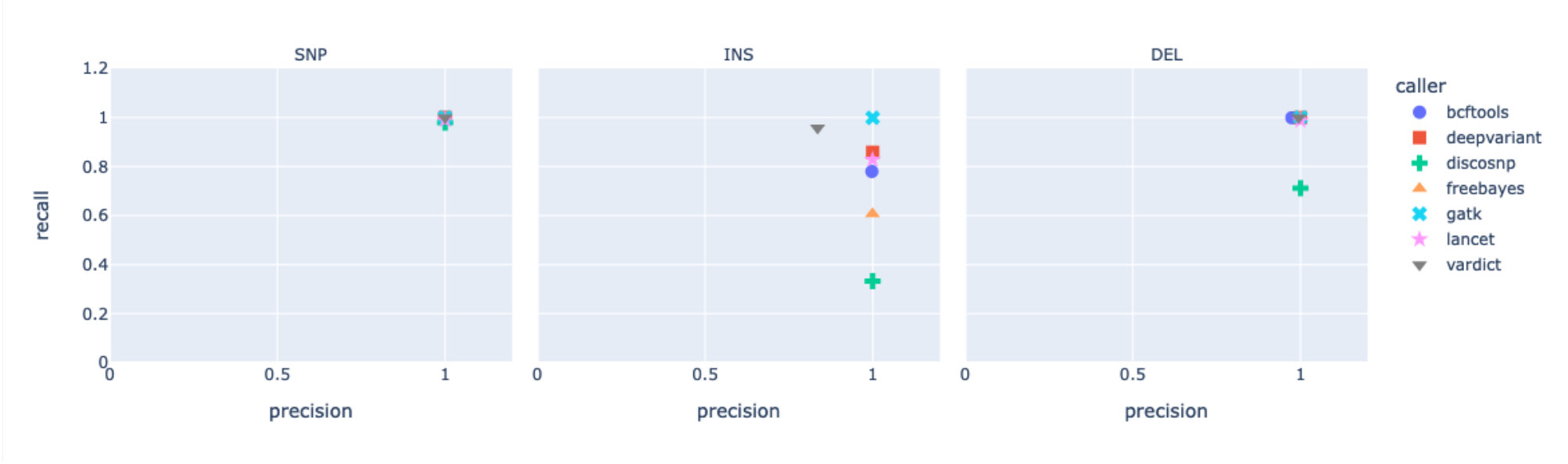
Recall vs. precision on synthetic datasets with SNP-only (left), INS-only (middle), and DEL-only mutations (right). Each dataset comprised 60 mutated genomes (1 mutation/200 bases) and synthetic reads generated at 50X average read depth.

We also investigated the effects of varying read depths from 5X up to 100X on variant caller performance. Mutations were introduced at a density of one for every 1,000 bases, resulting in a total mutation count of 4,398 for the SNPs-only dataset (Fig. 2a); 4,315 for the insertions-only dataset (Fig. 2b); and 4,261 for the deletions-only dataset (Fig. 2c). Across all variant callers, recall and precision in the SNPs-only dataset were ≥ 95% and ≥ 99% respectively, at read depths of 20X and higher (Fig. 2a).

**Figure 2.**
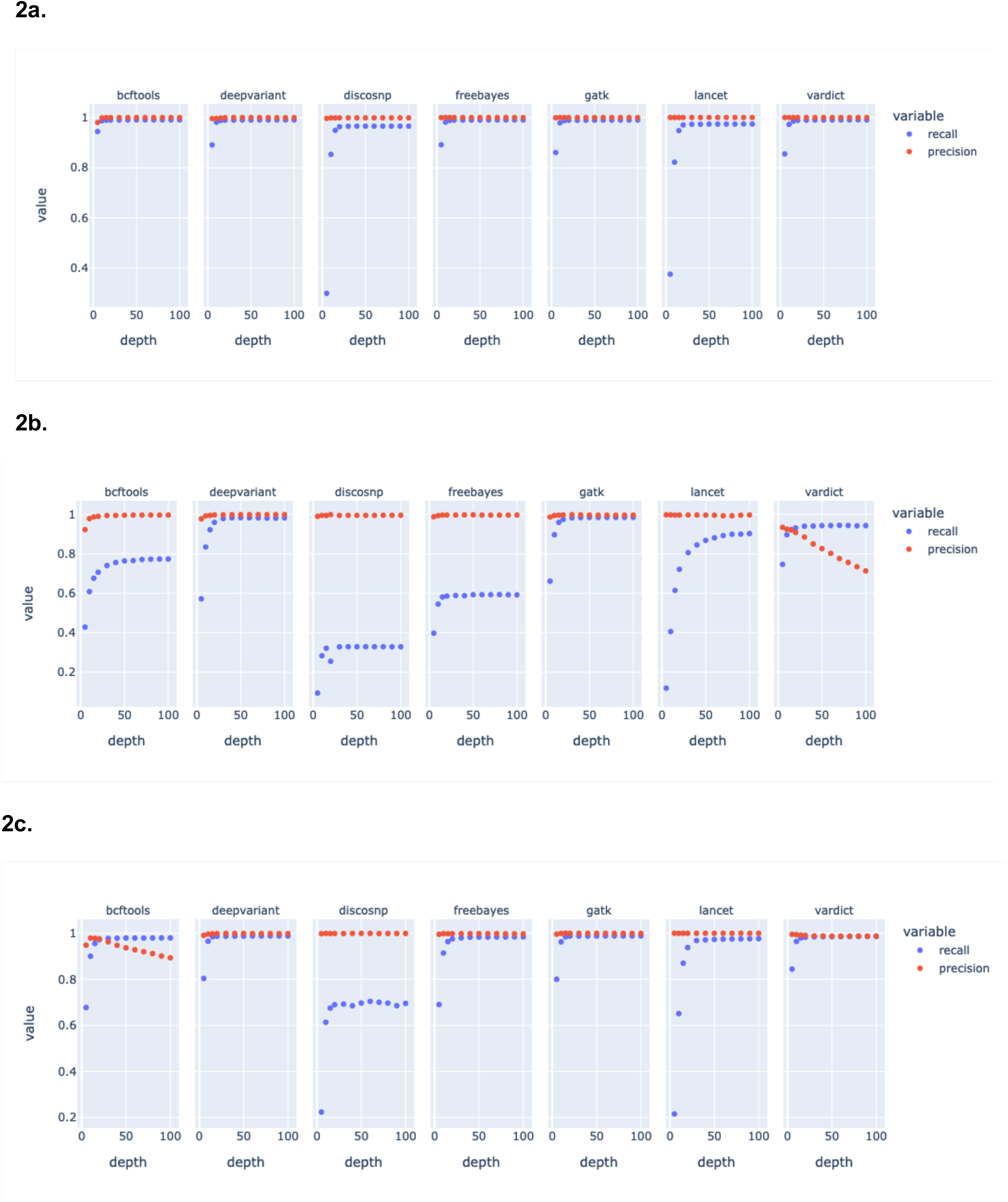
Recall and precision on synthetic datasets with 1 mutation / 1,000 bases per simulation, at varying read depths for the a. SNP-only, b. INS-only, and c. DEL-only datasets.

For the insertions-only dataset, however, recall at 20X ranged from 25.5% (DiscoSNP) to 97.5% (GATK) (Fig. 2b). The influence of read depth on recall also varied widely by variant caller, with the point of rapid decline in performance at 70X coverage for Lancet (89% recall) and lowest of 15X for FreeBayes (58% recall) (Fig. 2b). Precision of the variant callers on the insertions-only dataset at 20X was similar to the SNPs-only dataset at ≥ 99%, except for VarDict at 90.8%. Interestingly, VarDict’s precision was inversely correlated to read depth, steadily decreasing from 93% at 5X to 71% at 100X.

All the variant callers performed slightly better on the deletions-only dataset than on the insertions-only dataset at 20X read depth (Fig. 2c), with recall ranging from 68.9% (DiscoSNP) - 98.7% (GATK) and precision ranging from 97.55% (DiscoSNP) - 99.98% (GATK and Lancet). VarDict’s precision only declined slightly between 5X and 30X to 98% on the deletions-only dataset; however, precision of bcftools dropped from 97.9% to 89.3% between 10X and 100X. Three types of insertions were introduced into the insertions-only dataset referenced in Fig. 2b: duplications, inversions, and random sequences, at a ratio of 2:1:1. In order to further investigate the features of insertions that determine recall for each variant caller, we subdivided the same insertions-only dataset by the types of insertions introduced (Fig. 3): duplications-only (2,220 total mutations), inversions-only (1,016), and random sequences only (1,079). Median insertion lengths were 26 bp (mean 25.7, s.d. 14.4), 25 bp (mean 25.6, s.d. 14.3), and 24 bp (mean 24.8, s.d. 14.6) for duplications, inversions, and random sequences respectively. At read depths of 20X and higher, recall did not significantly differ between duplications, inversions, and random sequences for DeepVariant (respectively 93.6%, 98.1%, and 98.7%, at 20X) and GATK (96.7%, 98.1%, and 98.4%). Recall of duplications were the lowest of the three insertion types however, for DiscoSNP (max. 25.3% at ≥ 40X), FreeBayes (max. 47.7% at 40X), and bcftools (max. 67.9% at 90X), while recall of inversions were lowest for Lancet (max. 69.5% at 100X) and VarDict (max. 87.9% at 70X and 100X).

**Figure 3.**
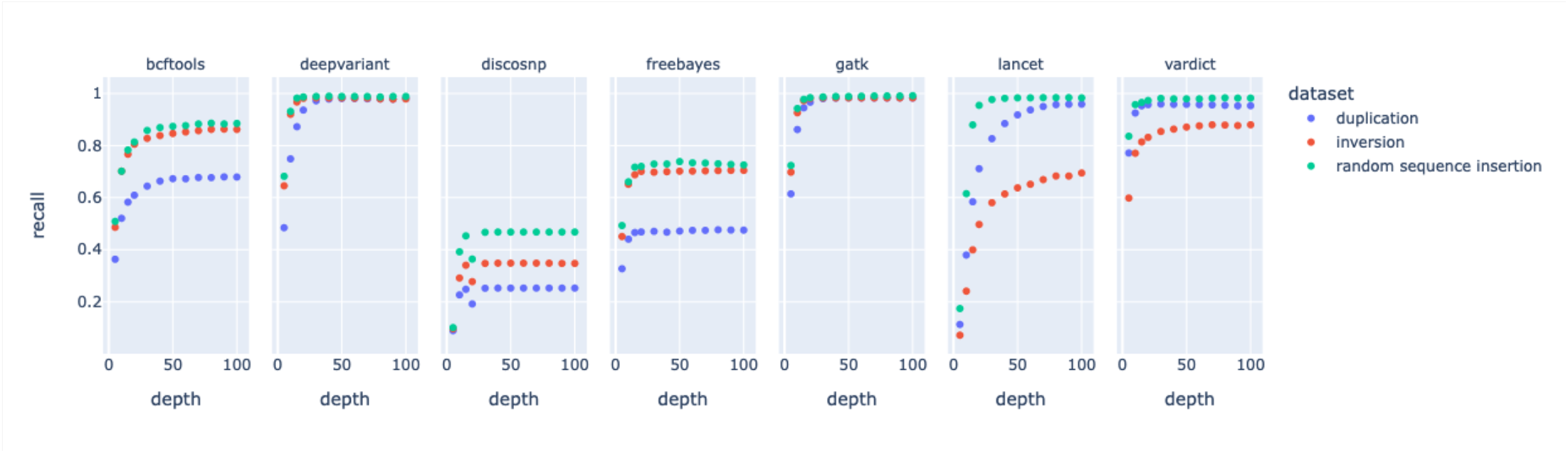
Recall values for insertions consisting of duplications, inversions, and random sequences. Precision was not calculated since false positive calls did not have information on the type of insertions that were identified by the variant callers.

Finally, we analyzed datasets consisting of mutations pooled from 60 simulated genomes, each with mixed mutation types introduced at a density of one every 200 bases (approximately 1.2 million mutations per dataset). Using this approach, we were able to generate a large number of observations without over-saturating individual genome locations. The first dataset comprised 50% SNPs and 25% each of insertions and deletions, while the second dataset consisted of 85% SNPs and 7.5% each of insertions and deletions (Fig. 4); insertions in both datasets comprised 50% duplications, and 50% inversions or random sequences. Median indel length was 25 bp (mean 25.5, s.d. 14.4). Recall and precision on the 50%-SNPs dataset ranged from 74.79% (DiscoSNP) - 99.95% (GATK), and 95.26% (VarDict) - 99.97% (FreeBayes, GATK, and Lancet), respectively. Recall and precision on the 85%-SNPs dataset meanwhile, ranged from 87.82% (DiscoSNP) - 99.98% (GATK), and 96.67% (DiscoSNP) - 99.99% (DeepVariant, FreeBayes, GATK, and Lancet), respectively.

**Figure 4.**
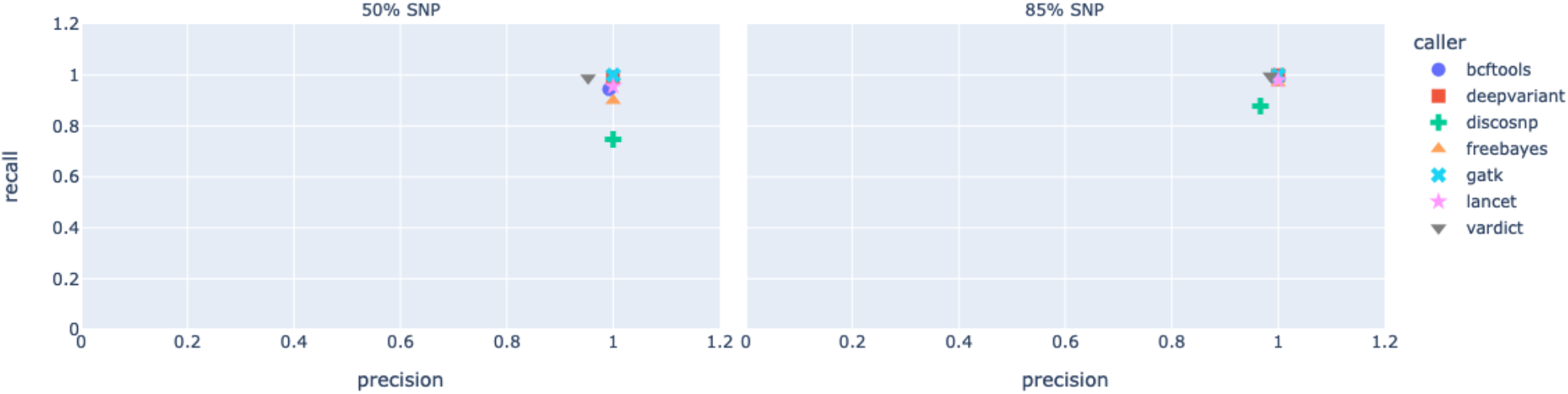
Precision vs. recall on synthetic datasets with mixed variants. Left: 50% SNPs, 25% insertions, 25% deletions; Right: 85% SNPs, 7.5% insertions, 7.5% deletions.

## Discussion

Identifying single nucleotide polymorphisms from whole genome sequencing short reads has become a relatively straightforward task that most popular variant calling software and pipelines can reliably perform, especially with high quality sample preparation and reference databases. More recent advancements in identifying SNPs focus on either calling with limited signal information (e.g., low-frequency variants), or reducing computational burden. Accurately identifying indels and structural variants, however, remains a challenge. Methods for indel detection generally employ gapped alignment, split reads, *de novo* assembly, or a combination thereof (7). Gapped alignment methods detect small indels that are contained within the length of a read; split-read methods identify medium-length indels (10 bp – 1 kb) at the cost of missing low-frequency indels, while *de novo* assembly methods can identify larger indels, albeit by utilizing significant computational resources (7).

Given that identifying variants is an integral part of clinical genomics, variant callers and other software used as part of the bioinformatics workflow are often validated and benchmarked against human reference datasets (e.g., (6–8)). Variant calling in the context of resolving clinical infections or tracking infectious disease transmission and epidemiology also relies on the same bioinformatics tools; therefore, these tools also need to be tested against microbial datasets. Recent efforts to validate widely used bioinformatics tools with microbial data include testing SNP-calling assemblies and pipelines against increasingly divergent bacterial reference genomes (1, 4), and combining whole genome sequencing and traditional molecular biology assays to validate a SNP-calling bioinformatics workflow (9). Steglich and Nübel (10) tested four indel callers against 1- to 2,321-bp indels introduced into a reference *Clostridium difficile* bacterial genome *in silico*, and found that while gapped alignment (e.g., as implemented in FreeBayes) was suitable for indels ≤ 29 bp, a combination of methods as employed by the ScanIndel framework (7) was best for indels > 29 bp.

In the present study, we synthetically generated variant whole genomes based on the *M. tuberculosis* H37Rv reference genome to test the performance of seven variant callers that are designed to identify both SNPs and short indels. The widely used ‘bcftools mpileup’ and ‘bcftools call’, which evolved from ‘samtools mpileup’ and ‘bcftools view’, applies a multiallelic variant-calling model to identify SNPs and indels by statistically inferring allele and haplotype frequencies based on “pileups” of reads aligned to a reference genome (11, 12). FreeBayes also models multiallelic loci within a Bayesian statistical framework based on aligned reads in order to detect variation across inferred haplotypes (13). DeepVariant meanwhile, takes advantage of images of read pileups and known true diploid genotypes to train a convolutional neural network model to subsequently identify new candidate SNP and indel variants (14).

Instead of strictly utilizing genomic position-based pileups, GATK HaplotypeCaller uses a consensus of reads at genomic regions of interest to assemble theoretical haplotypes from deBruijn-like graphs (15–17). VarDict similarly relies on aligned reads, then uses soft-clipped reads for local realignment to estimate indel allele frequencies (18). The somatic variant caller Lancet also locally assembles mapped reads, then decomposes them into k-mers to construct deBruijn graphs that are color-labeled to differentiate tumor versus normal samples (19). DiscoSNP meanwhile eschews aligned reads completely, instead constructing deBruijn graphs of k-mers from raw read sets to detect SNPs and indels (20).

While all tested variant callers generally excelled at identifying SNPs, the two variant callers that were best at recalling insertions (Fig. 1b), GATK HaplotypeCaller and VarDict, both scrutinize genomic regions with discrepancies such as base mismatches and high quality soft-clipped bases to identify indels (17, 18). This also suggests an explanation for DiscoSnp’s poor recall, since local alignment context is lost when only k-mers of raw reads are used for variant identification. GATK HaplotypeCaller and VarDict’s high recall corresponds with the performance of the ScanIndel framework – the latter also recovers soft-clipped reads for realignment before final variant calling, which helps to identify medium-sized insertions and large deletions (7). Interestingly, higher coverage depth up to 100X and lower mutation density caused VarDict to falsely call more insertions than GATK HaplotypeCaller did (Fig. 2b), by up to three orders of magnitude. Lai et al. (18) describes VarDict’s strength in detecting deletions and complex variants that include deletions, which are supported by our observations in the deletions-only dataset. The deletions-only dataset also posed less of a challenge to the other tested variant callers, presumably because deletion breakpoints are simpler to identify by the lack of reads or read segments.

The variant callers surveyed in this study employ different strategies that ultimately attempt to deal with the problem of aligning reads that do not match the reference sequence. Our data indicate that algorithms that take advantage of local genomic context, especially when there are significantly mismatched or soft-clipped bases, perform better than those that do not; however, the specific local realignment strategies adopted by the variant callers matter in differentiating their respective performances on insertions versus deletions.

## Methods

### Variant Simulation

A locally developed Python script was used to introduce SNPs, insertions, and/or deletions into the *M. tuberculosis* H37Rv whole genome sequence (GenBank accession NC_000962.3) to generate novel variant genomes *in silico*. All pipeline associated code are publicly available at https://github.com/molmicdx/mtb-pipeline.

We generated multiple replicates of synthetic variant genomes comprising SNPs only, insertions only, or deletions only to test the performance of variant callers on each type of mutation. In order to obtain adequate numbers of mutations for statistical inference with a low probability of introducing adjacent mutations, we performed 60 replicates with a mutation density of 1 mutation every 200 bases; since H37Rv has a genome size of 4,411,532 bp, this gave us approximately 20,000 mutations per variant genome. Indel length was allowed to range between 1 and 50 bases, resulting in median indel length of 25 bases (mean 25.5, s.d. 14.4). We also generated synthetic variant genomes that comprised SNPs only, insertions only, or deletions only, with a mutation density of 1 every 1,000 bases (approximately 4,000 mutations per variant genome) to model genetic distance in the same order of magnitude as those between MTBC strains, which have average nucleotide identities of greater than 99% (21). Validation of introduced mutations as done by aligning reference and selected variant genomes in Mauve (22), followed by manually inspecting the sequences at the mutated coordinates.

### Read Simulation, Processing, and Mapping

Synthetic paired-end reads were generated from the H37Rv reference genome and simulated variant genomes with ART, with read lengths set at 150 bp (mean fragment length was 200 bp, s.d. 10 bp). The average read coverage for the replicate datasets with density of 1 mutation / 200 bases was set to 50X, while reads generated for the datasets with 1 mutation / 1,000 bases had average coverage of 100X. The latter datasets were then subsampled to average coverages of 90X, 80X, 70X, 60X, 50X, 40X, 30X, 20X, 15X, 10X, and 5X to investigate the effect of read depth on variant calling. Instead of the default base quality profile model, we provided our own, based on in-house sequencing of *M. marinum*. The synthetic reads were trimmed with cutadapt, retaining those with minimum read quality of 5 and minimum length of 20 bp. We also used cutadapt to detect and trim Illumina adaptor sequences.

Pre-processed reads were then mapped to the H37Rv reference genome with the ‘bwa mem’ algorithm. The resulting BAM files were coordinate-sorted and indexed with samtools. Duplicated reads and poorly-mapping reads (MAPQ < 10) were removed using GATK MarkDuplicates (Picard) and ‘samtools view’, respectively.

### Variant Calling

The final BAM files were used as input for six different variant callers: bcftools, DeepVariant, FreeBayes, GATK HaplotypeCaller, Lancet, and VarDict. We also tested a seventh variant caller, DiscoSNP, which is a reference-free method that does not use BAM files as input; instead, DiscoSNP uses raw read sets in kmer-based deBruijn graph analysis to identify variants. While the somatic variant caller Lancet also uses kmer-based deBruijn graphs, Lancet decomposes mapped reads from BAM files. In order to take advantage of Lancet’s joint analysis of tumor and normal samples, we simulated and mapped reads from the H37Rv reference genome to use as the normal reference reads; see *Read Simulation, Processing, and Mapping* above.

Where the options were available, ploidy was set to one (bcftools, FreeBayes, GATK), and minimum alternate allele fraction set to 0.2 (FreeBayes, VarDict, Lancet) to reflect expected variant frequencies in haploid genomes. Other than ploidy and alternate allele fraction, default parameters were used in all variant callers; we assumed the multiallelic variant-calling model as the default setting for ‘bcftools call’.

### Variant Normalization and VCF Filtering

Since each variant caller can have different default information reported in their output VCFs, we post-processed all VCFs so that variant sequences and genomic coordinates could be accurately compared to determine true/false calls for calculation of precision and recall. True positive variant calls are those with the same sequence and genomic coordinates as simulated mutations. Variant calls that differ from the simulated mutations in sequence or coordinates are considered false positive, while false negative calls are simulated mutations that are not identified by the variant caller. Precision (or positive predictive value) is then calculated as the ratio of total true positive calls to the sum of true positive and false positive calls, while recall (sensitivity) is the ratio of total true positive calls to the sum of true positive and false negative calls.

Variant normalization refers to both left-aligning the variant coordinates with respect to the reference genome, and parsimoniously representing the variant sequence. GATK LeftAlignAndTrimVariants was used to normalize variant representations in all VCF files. Since most variant callers report diploid genotypes, we decided that heterozygous alternate alleles should be counted as valid variant calls as well, and only filtered out homozygous reference alleles (e.g. GT = 0/0) from VCFs that reported them.

